# Serum response factor reduces gene expression noise and confers cell state stability

**DOI:** 10.1101/2022.08.04.502808

**Authors:** Jian Zhang, Xiao Hu, Qiao Wu, Shangqin Guo

## Abstract

The role of serum response factor (Srf), a central mediator of actin dynamics and mechanical signaling, in cell identity regulation is debated to be either a cell identity stabilizer or destabilizer. We thus investigated the role of Srf in cell fate stability using mouse pluripotent stem cells, one of the very few cell types that can tolerate null *Srf*. Despite the fact that serum-containing cultures yield heterogeneous gene expression, deletion of *Srf* in mouse pluripotent stem cells leads to further exacerbated cell state heterogeneity. The exaggerated heterogeneity is not only detectible as increased lineage priming, but also as the 2C-like cell state. Thus, pluripotent cells explore more variety of cellular states in both directions of development surrounding naïve pluripotency, a behavior that is constrained by Srf. These results support that Srf functions as a cell state stabilizer, providing rationale for its functional perturbation in cell fate engineering and pathological intervention.

## Introduction

Robust cell state stability is required for effective execution of specific biochemical and biophysical functions, loss of which underlies degeneration ^1–4^. On the other hand, noisy cell states also contribute to plasticity, enabling the emergence of new cell identity ^5^ required for development and homeostasis throughout life. Consistent with this notion, cell-to-cell heterogeneity is known to impart development robustness ^6^ and culture conditions that reduce pluripotency gene expression noise also compromises multipotency ^7^. Pathological cell states, such as malignancy, could be viewed as cells in protracted state excursions or cells entrapped in a proliferative state ^8^. Organismal well-being rests on a delicate balance between cell state plasticity and stability. A particularly relevant biological context in which cellular heterogeneity is implicated is aging: increased cellular heterogeneity is one of the hallmarks of aging ^9,10^, coupled with loss of cell identity ^11^. Decreased SRF activity has been seen in multiple aging systems ^12^ and that instigators of SRF signaling has recently been shown to rejuvenate oligodendrogenesis in aged mice ^13^.

SRF is a ubiquitously expressed transcription factor (TF), responding to many extracellular signals to control the expression of two large categories of genes. One is components of the actomyosin cytoskeleton, in complex with the myocardin family of transcriptional coactivators, MKL1/2. These genes are involved in adhesion and motility ^14,15^ and underlies cell size and morphology. The other major group of SRF target genes is the ‘immediate-early’ genes, such as *Jun, Fos, Egr1*, whose expression is rapidly activated by mitogenic stimuli ^16,17^. Strikingly, cell size/morphology and immediate early gene expression are the top features highly predictive of gene expression heterogeneity ^18^. Furthermore, SRF is a major mediator of circadian oscillation ^19,20^. Therefore, it is conceivable that the cell-to-cell variability in gene expression is significantly contributed by SRF, whose loss-of-function could confer a less variable gene expression state. However, SRF inactivation is generally not tolerated in cells post E6.5 ^21–24^, which either leads to cell or animal demise or compensatory cells escaping Cre-mediated conditional allele excision, confounding the assessment of cellular heterogeneity. Unlike somatic cells, SRF is dispensable for pluripotent stem cells for both human ^25^ and mouse ^17^. We previously reported the derivation of *Srf* null iPSCs ^26^, providing the cellular system to examine the role of SRF in controlling cell state heterogeneity in pluripotent stem cells.

Mouse pluripotent stem cells can interconvert between several related states ^27^. For example, they can be maintained in the naive pluripotent state (ICM-like) or primed pluripotent state (epiblast-like) depending on culture conditions ^28,29^, with the latter being a developmentally more advanced stage than the naïve state. Signaling pathways, such as the MAPK cascade and retinoic acid signaling, induces the exit from naïve pluripotency and committing to specific lineages. Preceding the pluripotent ICM cells, the zygote and 2-cell (2C) stage blastomeres are totipotent ^30^. The 2C-like cells can arise spontaneously in mESC cultures at low frequency (~ 0.5%) ^30^. Similar to 2-cell-stage embryos, 2C-like cells reactivate 2-cell-specific transcripts including *Zscan4, Eif1a-like, Gm6763*, major satellites (MajSat) repeats and murine endogenous retrovirus with leucine tRNA primer (*MuERV-L*, also known as *MERVL* and *Erv4*)^30–32^. As each cell state is associated with different developmental potential, the molecular control of the interconversion between these cell states have been intensely studied. For example, depletion of the p150 or the p60 subunits of chromatin assembly factor-1 (CAF-1), or expression of *Dux* in mESCs promotes the emergence of 2C-like-cells ^33–35^. Taken together, exiting naïve pluripotency to enter the 2C-like state or lineage priming are developmentally divergent and are under distinct regulatory mechanisms. Here, we report the surprising finding that these divergent cell states could both arise following the loss of a single gene, *Srf*

## Results

### *Srf* null iPSCs display heightened cell state heterogeneity

*Srf* null (*Srf^Δ/Δ^* iPSCs were generated by reprogramming myeloid progenitors of adult reprogrammable mouse, followed by Cre-mediated excision of a conditional *Srf* allele, as we described before ^26^. Consistent with the previous reports that *Srf* null mESCs remain pluripotent, these *Srf* null iPSCs can be propagated in mESC culture conditions, display reduced cortical actins and maintain highly accessible chromatin ^26^. To explore whether and how SRF controls cell state heterogeneity, *Srf* null (KO) or wild type (WT) iPSCs cultured in serum containing medium were subject to single-cell RNA sequencing (scRNA-seq). After quality filtering and normalization, 857 cells for WT and 960 cells for *Srf* KO were retained. The mean and median numbers of detected genes per cell were 3915 and 4135 for KO and 4114 and 4275 for WT, respectively (Supplementary Figure 1). Using the 3,000 most variable genes, Manifold Approximation and Projection (UMAP) dimensional reduction identified two major cell clusters (Fig. 1A): a major cluster encompasses most cells in both genotypes with a smaller cluster limited to *Srf* KO cells (Fig. 1A). *Srf* is not detected in SRF KO cells, confirming the inactivated status of the gene. Even in the WT population, only ~20% (163 cells) displayed detectible *Srf* expression, indicating that WT iPSCs naturally display a spectrum of *Srf* levels (Fig. 1B). In addition to *Srf* itself, the expression of an Srf target gene *Actb* mirrors that of *Srf*, consistent with its greatly reduced transcriptional activity. In contrast, the expression of core pluripotency genes including *Pou5f1* (*Oct4*) and *Nanog* are similar in both genotypes (Fig. 1B), confirming the pluripotent nature of both WT and KO cells.

**Figure 1.**
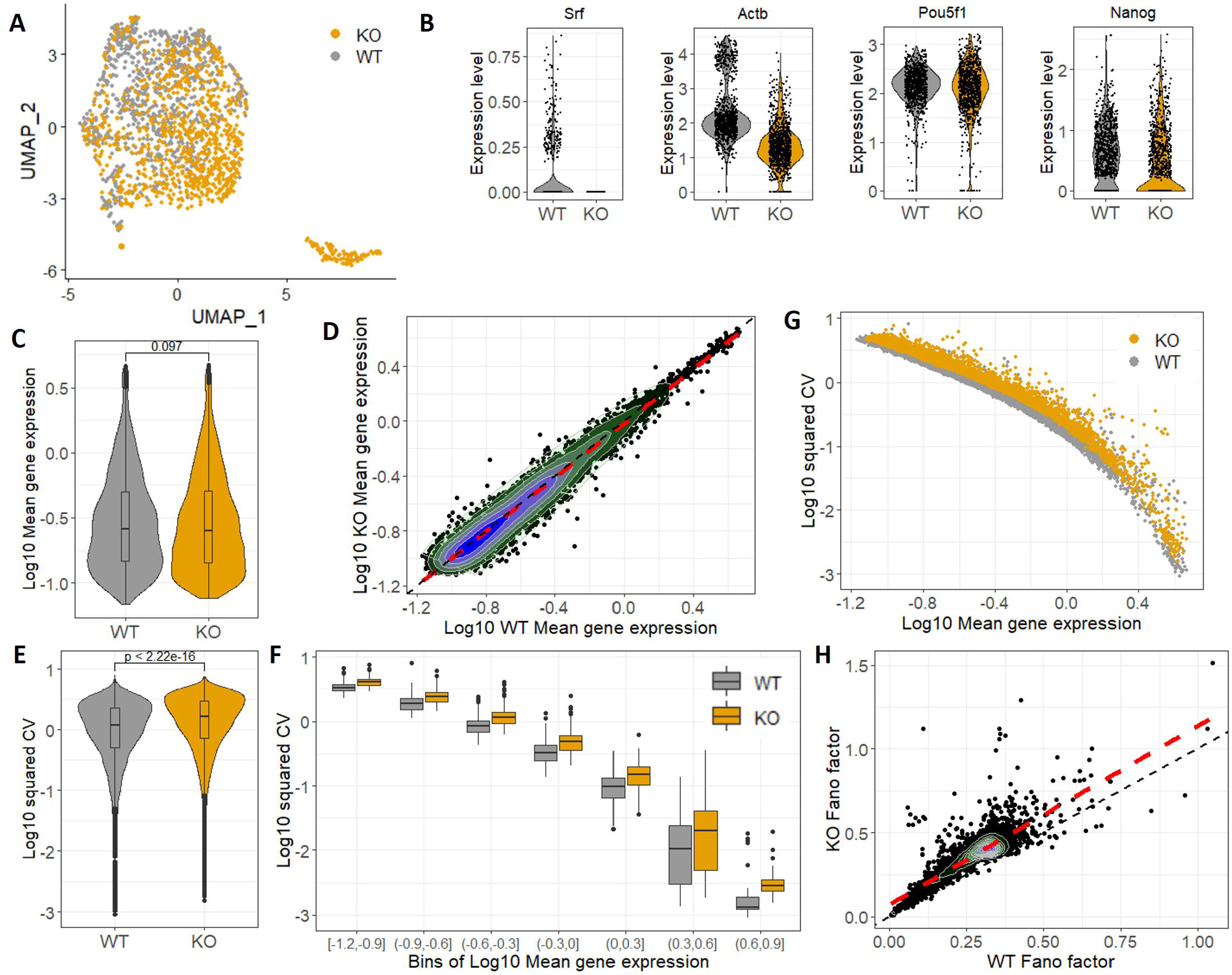
Loss of *Srf* increases transcriptional heterogeneity of mouse iPSCs. (**A**) Projection of all cells onto two principal dimensions using UMAP based on the 3,000 most variable genes. Genotypes are distinguished by color. (**B**) Violin plots of select genes in *Srf* KO and WT iPSCs. Lower expression in *Srf* and *Actb* confirms the null status of *Srf* in KO cells. (**C**) Violin plots depicting the mean gene expression levels in *Srf* KO and WT iPSCs. Median and 95% CI of mean gene expression levels for the 5943 genes in the Robust gene set (Supplementary Tables 2) are shown. Wilcoxon rank test, p-value =0.097. (**D**) Dot plot depicting the mean gene expression of 5943 genes in *Srf* KO vs. WT iPSCs. (**E**) Violin plots depicting the Coefficient of Variation (σ^2^/μ^2^, CV^2^) of gene expression in *Srf* KO and WT iPSCs. Wilcoxon rank test, p-value < 2.22e-16. (**F**) Binned gene expression CV^2^ across the all gene expression averages for *Srf* KO and WT iPSCs. Wilcoxon rank test, p-value < 1e-5 for all bins. (**G**) Mean expression vs. CV^2^ for all 5943 genes. (**H**) Distribution of the Fano factor (σ^2^/μ) of all 5943 genes in *Srf* KO vs. WT iPSCs. Overlay of density contours reveals how center of mass (red dashed line) lies above the diagonal line, as shown for mean values in D and the black dashed lines.

We next assessed the transcriptome-wide cell-to-cell variation in gene expression levels (i.e. intercellular gene expression heterogeneity) in *Srf* KO and WT iPSCs. While the mean-expression levels for most genes exhibit minimal changes between KO and WT iPSCs (Fig. 1C,D), *Srf* KO iPSCs displayed higher cell-to-cell variability in transcript levels (i.e., transcript noise) for virtually all genes across the genome, as analyzed by CV^2^ (σ^2^/μ^2^) versus mean (Fig. 1E-G).). Higher CV is seen in KO cells independent of the gene expression levels (Fig 1G).

Additionally, we quantified transcript noise using the Fano factor (σ^2^/μ). Despite mean-expression levels exhibiting minimal changes (Fig. 1C,D), the Fano factor increased for > 90% of genes in *Srf* KO iPSCs (Fig. 1H). These results indicate that *Srf* KO iPSCs display a global increase in transcript noise with little change in mean levels for the great majority of the transcriptome. Importantly, the noisy cell state is not contributed by the small population unique to the KO cells, as most KO cells display higher CV^2^ and Fano factor (Fig. 1E-H). Thus, pluripotent stem cells exist in a state of heightened heterogeneity when *Srf* is inactivated.

### *Srf* KO iPSCs display increased expression of differentiation markers

To examine the cellular consequence of this gene expression heterogeneity, we sought to examine the potential emergence of cell states deviating or departing from the naïve pluripotent cell state. While scRNAseq data resolves cell state heterogeneity, the data is often sparse and may not capture the expression of genes that are present in low levels or transiently. Therefore, we performed bulk RNA-seq of parallel *Srf* KO and WT iPSCs. We identified 816 up-regulated and 247 down-regulated differentially expressed genes (DEGs) (Fig. 2A and Supplementary Tables 1). Gene set enrichment analysis (GSEA) indicated that epithelial-to-mesenchymal transition (EMT), inflammatory and interferon pathways were up-regulated in *Srf* KO iPSCs compared with WT controls (Fig. 2B). Many Srf target genes including *Actb*, *Actg1*, *Fos* and *Egr1* were substantially down-regulated, corroborating Srf loss of function (Fig. 2C).

**Figure 2.**
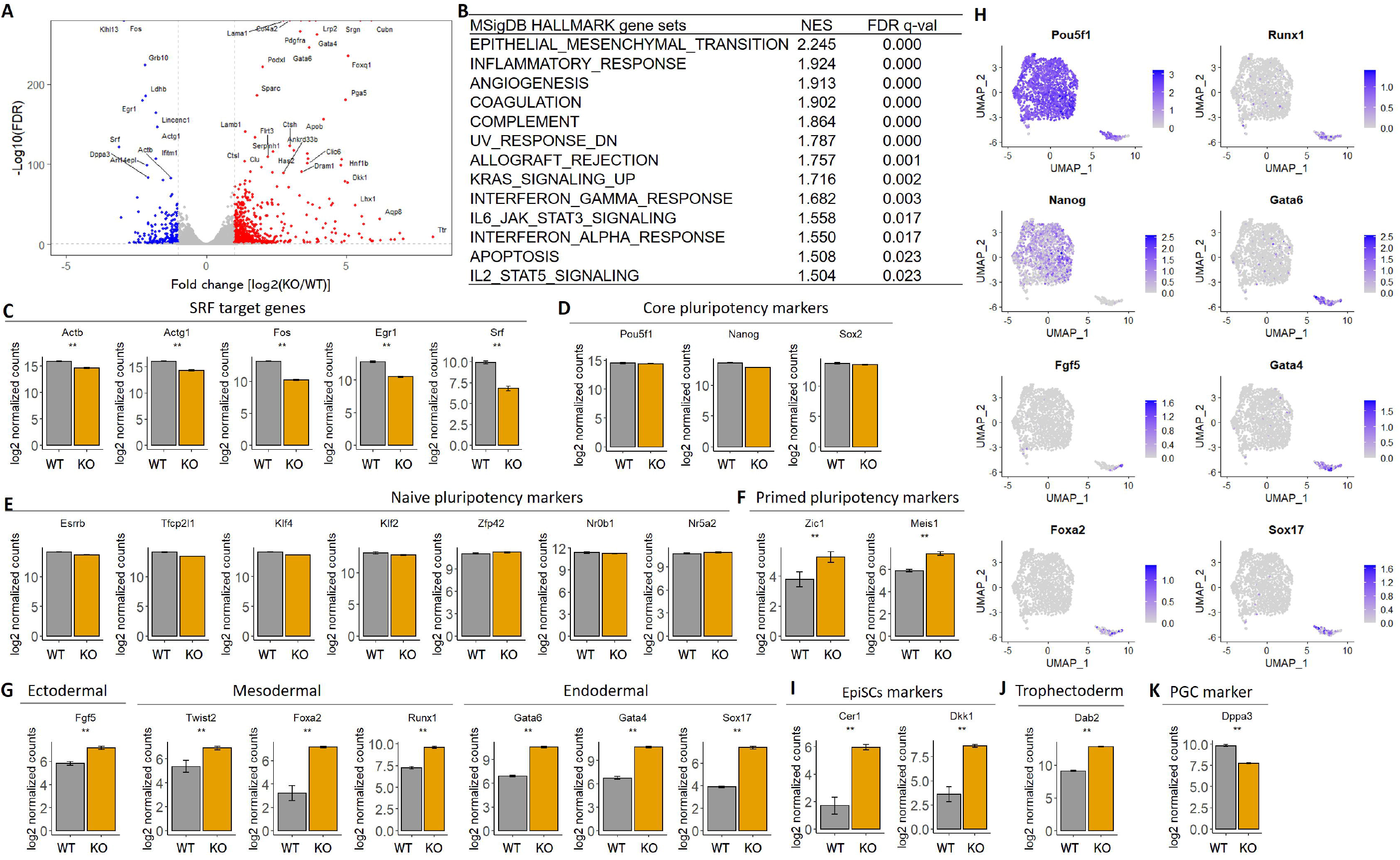
*Srf* KO iPSCs display increased expression of differentiation markers. **(A)** Volcano plot of differentially expressed genes (DEGs), defined as FDR-adjusted *P* value ≤0.05 and a log_2_ fold change of at least 1. The red and blue dots represent up-regulated and down-regulated DEGs in *Srf* KO iPSCs compared with WT controls. Shown are the top 40 most differentially expressed genes. (**B**) Upregulated genes in *Srf* KO iPSCs are enriched in multiple pathways, as detected by Gene Set Enrichment Analysis (GSEA) on MSigDB HALLMARK gene sets. **(C)** Expression of known *Srf* target genes in bulk RNA-seq of *Srf* KO and WT iPSCs. Lower expression of these genes confirm the reduced Srf function in the KO cells. (**D**) Expression of core pluripotency genes. (**E**) Expression of naive pluripotency marker genes. (**F**) Expression of primed pluripotency marker genes. (**G**) Expression of lineage specific genes, including ectodermal (*Fgf5*), mesodermal *(Twist2, Foxa2* and *Runx1)* and endodermal *(Gata6, Gata4* and *Sox17)* markers. (**H**) Expression of lineage genes on scRNA-seq UMAP. (**I**) Expression of epiblast stem cells (EpiSCs) marker genes. (**J**) Expression of trophectoderm marker genes. (**K**) Expression of primordial germ cell (PGC) marker genes. * |log2FC| > 1 && FDR < 0.05; ** |log2FC| > 1 && FDR < 0.01.

Consistent with the scRNAseq results, core pluripotency-associated genes such as *Pou5f1* (*Oct4*), *Nanog* and *Sox2* were unaffected (Fig. 2D). Expression of additional naïve pluripotency state genes including *Esrrb*, *Tfcp2l1*, *Klf4*, *Klf2*, *Zfp42*, *Nr0b1* and *Nr5a2* were also unaffected (Fig. 2E). These results further support that Srf is dispensable for naïve pluripotency. However, primed pluripotency genes such as *Zic1* and *Meis1* were significantly upregulated in *Srf* KO iPSCs (Fig. 2F).

To determine whether *Srf* KO iPSCs deviated more from pluripotency, we examined their expression of lineage-specifying transcription factors. *Srf* KO iPSCs displayed upregulation of markers of all three germ layers. These include the ectoderm lineage marker *Fgf5*; Mesoderm lineage markers *Twist2*, *Foxa2* and *Runx1*; Endoderm lineage markers *Gata6, Gata4* and *Sox17* (Fig. 2G). Moreover, these lineage specific genes were expressed in the small cluster specific to *Srf* KO in the scRNA-seq data (Fig. 2H). Lastly, epiblast stem cell (EpiSC) marker genes *Cer1* and *Dkk1* (Fig. 2I) and the trophectoderm marker gene *Dab2* (Fig. 2J) were significantly upregulated in *Srf* KO iPSCs. Taken together, when *Srf* activity is low or absent, pluripotent stem cells more frequently explore a differentiated/committed state. The only exception appears to be the primordial germ cell (PGC) lineage, whose marker gene *Dppa3* (*Stella*) was downregulated in *Srf* KO iPSCs (Fig. 2K), suggesting Srf’s role in supporting germ cell potential.

### *Srf* null iPSCs display increased expression of marker genes for 2C-like state

On the surface, the results above could be seen as *Srf* KO iPSCs are prone to differentiate, which is surprising given that *Srf* null embryos are defective in germ layer specification ^36^. An alternative interpretation is that *Srf* KO iPSCs display generally reduced cell state stability. In this scenario, one would expect to detect the emergence of other cell states, including those that is opposite to the developmental direction. One well known auxiliary cell state that pluripotent cells fluctuate in and out is the 2C-like cell state. Therefore, we compared our data to 2C-like embryonic stem cells (MERVL^+^/*Zscan4*^+^, *Zscan4*^+^) ^37–39^ by GSEA. The top of the *Srf* KO transcriptome was significantly enriched in the 2C-specific genes (2C-up) both in *Zscan4^+^* and MERVL^+^/*Zscan4*^+^ cells (Fig. 3A and B). *Srf* inactivation resulted in strong upregulation of the 2C-specific genes including *Zscan4* genes (*Zscan4a*, *Zscan4b*, *Zscan4c*, *Zscan4d*, and *Zscan4e*) and *Eif1a-like* genes (*Gm5662*, *Gm2022*, *Gm4027*, *BB287469*, *Gm2016*, *Gm21319* and *Gm8300*), *Dux*, *Zfp352*, *Usp17lb*, *Tcstv1*, *Tcstv3*, major satellites (MajSat), LINE1 and MERVL (Fig. 3C). These results indicate that deletion of *Srf* induces the emergence of 2C-like cells.

**Figure 3.**
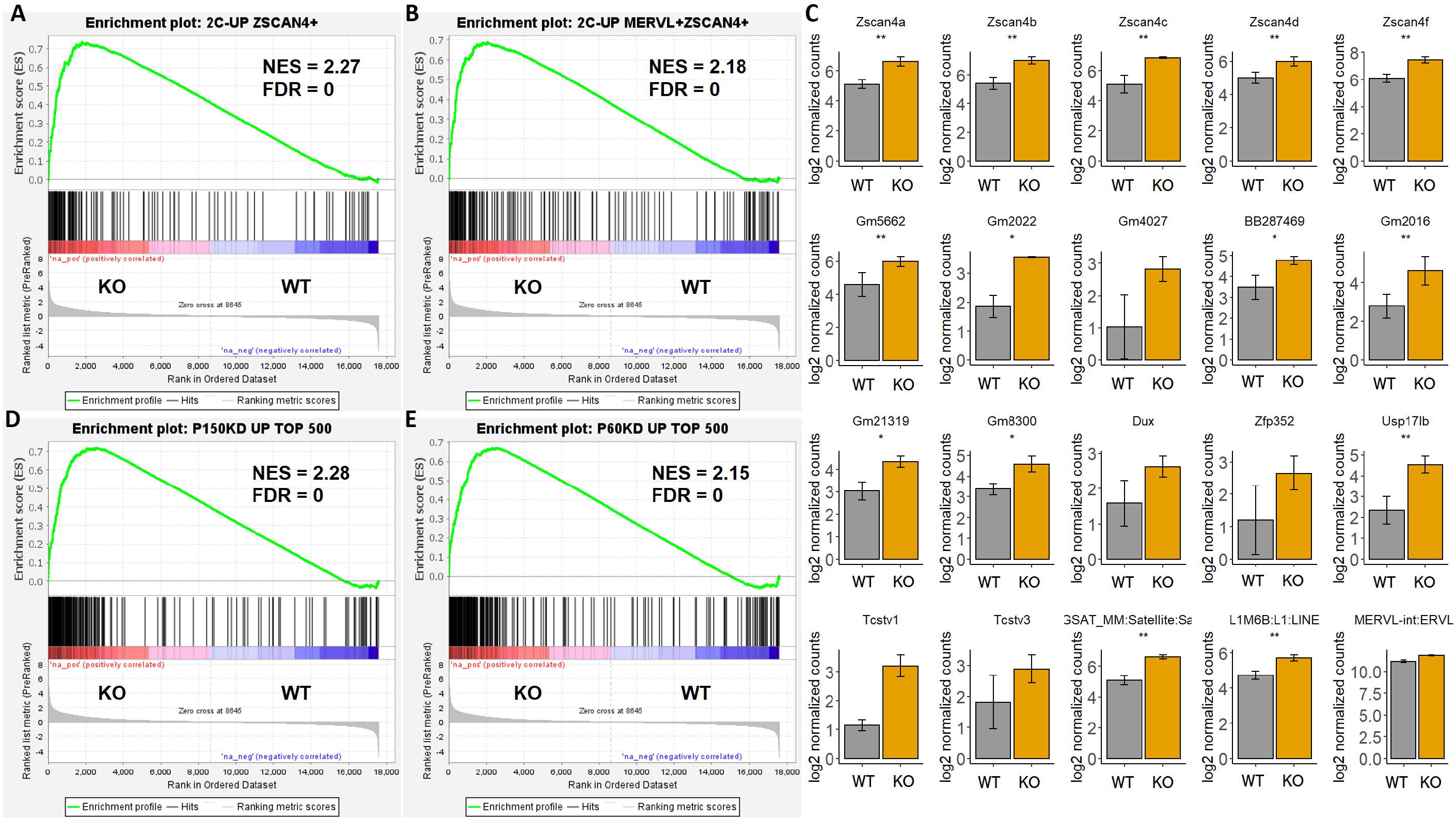
*Srf* KO iPSCs display increased expression of marker genes for 2C-like state. **(A)** GSEA of the 229 upregulated genes in the *Zscan4*^+^ 2C-like cells ^39^. The x axis shows the log2(fold change)-ranked *Srf* KO transcriptome. The enrichment score increases when a gene in the *Srf* KO transcriptome is also present in the 2C-like gene set, and a black vertical bar is drawn at bottom; the enrichment score decreases when a gene is absent in the 2C-like gene set. The p value was empirically determined on the basis of 1,000 permutations of ranked gene lists. **(B)** GSEA of the 310 upregulated genes in the MERVL^+^/*Zscan4*^+^ 2C-like cells ^39^. **(C)** Expression of a panel of 2C-like genes between *Srf* KO and WT iPSCs. * |log2FC| > 1 && FDR < 0.05; ** |log2FC| > 1 && FDR < 0.01. **(D)** GSEA of the top 500 most upregulated genes in p150 knockdown (KD) mESCs. **(E)** GSEA of the top 500 most upregulated genes in p60 KD mESCs.

In addition to natural fluctuation, 2C-like cells can be induced following specific perturbation. Ishiuchi et al reported that depletion of either the p150 or the p60 subunits of chromatin assembly factor-1 (CAF-1) in ES cells increases the population of 2C-like-cells ^33^. Therefore, we compared our *Srf* KO transcriptome with that of p150 or p60 knockdown (KD). Strikingly, we observed significant overlap between the transcriptomes following *Srf* loss-of-function with those following p150 or p60 knockdown (Supplementary Figure 2A and B; p-value < 2.2e-16 and p-value = 4.298e-16 for p150 and p60 RNAi, respectively, by two-sided Fisher’s exact test). Moreover, GSEA indicates that the most up-regulated genes in *Srf* KO iPSCs are enriched in the most up-regulated genes in p150 or p60 KD (Fig. 3D and E). These results demonstrate that *Srf* KO iPSCs and *Caf-1* KD cells share substantial overlapping gene-expression patterns.

Taken together, *Srf* loss of function in pluripotent stem cells increases the likelihood of otherwise rare or transient cell states, including the 2C-like cells and various lineage marker positive cells. These results collectively argue that Srf functions as a cell fate stabilizer. In its absence, gene expression becomes noisier.

## Discussion

Our results here suggest that mouse pluripotent stem cells, at least in serum containing culture conditions supporting naïve pluripotency, could depart from this main state and enter other cell states in both developmental directions. This cell state excursion is constrained, at least partly by the ubiquitously expressed transcription factor *Srf.* Its role in stabilizing the naïve pluripotent cell state extends our previous work in identifying the actin-MKL1/SRF pathway in stabilizing somatic cell fate ^26^. In the case of somatic cell reprogramming driven by the Yamanaka factors, the activity of actin-MKL1/SRF pathway needs to be sufficiently weakened to allow the entry into pluripotency. Together, our results lend further support that Srf is part of the actin-MKL1/SRF pathway contributing to cell fate stability. With Srf appreciated as a cell fate stabilizer, its function in somatic and pluripotent stem cells can be understood in one unified model (Fig. 4). Our results offer a potential molecular explanation why certain small molecules with diverse targets or mechanisms, such as ROCK inhibitors and Arp2/3 inhibitors, are potent cell fate modulators ^26,40,41^. On the other hand, once the desired cell fate is attained and needs to be propagated, further attenuation in Srf could be counterproductive. These insights contrasts that of a recent study suggesting that Srf destabilizes somatic cell identity ^42^.

**Figure 4.**
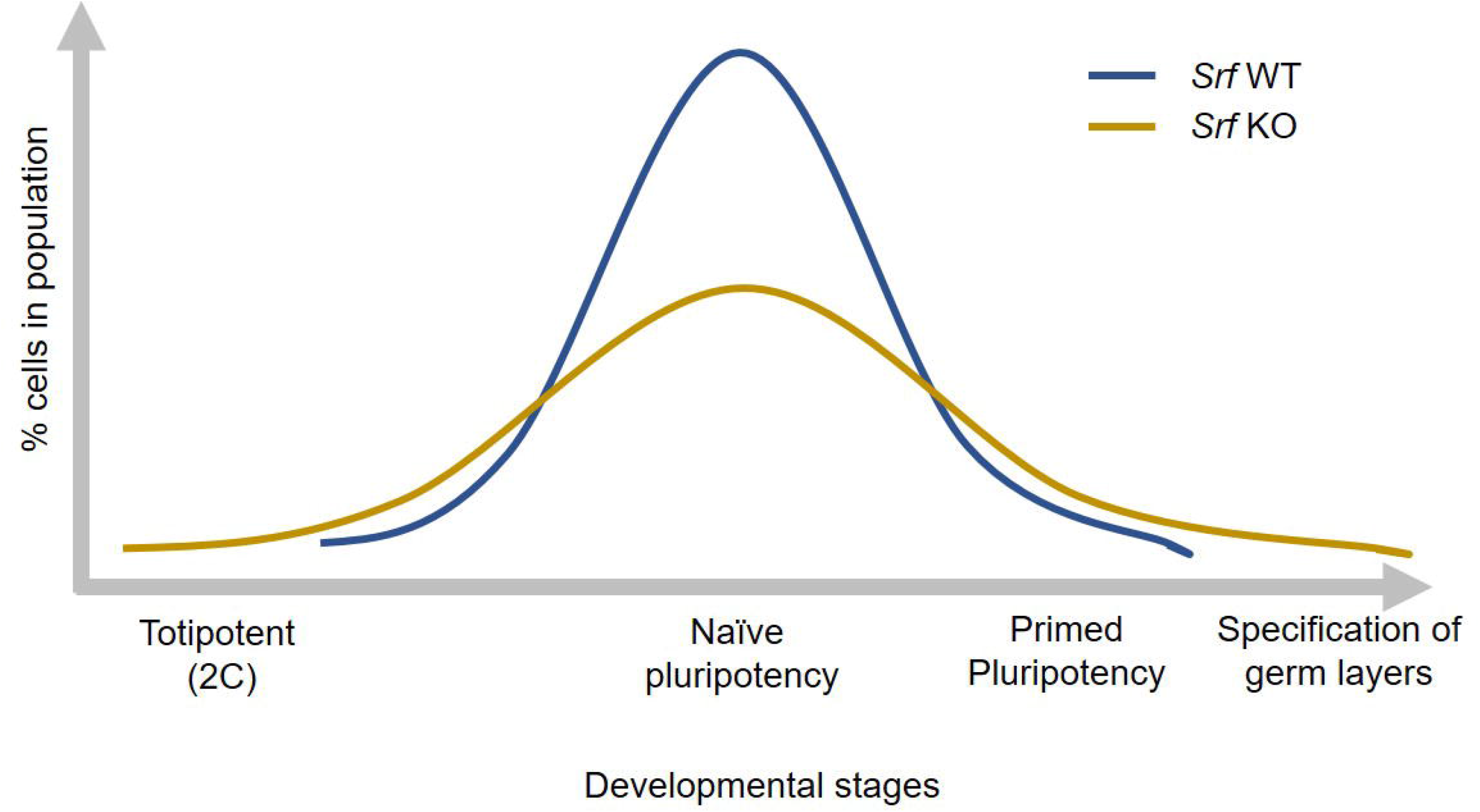
Working model of Srf conferring cell state stability in pluripotent stem cells. Pluripotency state, at least when cultured in serum containing medium, is heterogeneous containing a spectrum of cell states arising spontaneously (blue line, WT). The spread of these cell states is constrained by *Srf* (yellow line, KO). Deleting *Srf* allows more cells to appear in the tail regions in both directions of development.

As a ubiquitously expressed transcription factor, how Srf regulates cell state heterogeneity remains to be identified. A recent report identified that DNA repair pathway regulates gene expression noise and cell fate plasticity ^5^. Since human pluripotent stem cells under prolonged culture often display DNA replication stress and chromosomal abnormalities, a process that has been linked to reduced SRF expression ^25^, SRF may reduce gene expression noise by promoting DNA damage repair. Consistent with this possibility, our GSEA results did indicate that inflammatory and apoptotic pathways to be upregulated in *Srf* KO iPSCs (Fig. 2B). Beyond pluripotent stem cells, *Srf* loss-of-function has been reported in several human diseases via alternative splicing or caspase mediated cleavage among others, often yielding a dominant negative form of the transcription factor ^43–46^. Our results could provide insights into the various pathological conditions.

## Methods

### Bulk RNA sequencing (RNA-seq) and data analysis

Mouse iPSCs with *Srf^Δ/Δ^* (KO) and *Srf^f/f^* (WT) were generated and cultured in serum containing medium as we reported before ^26^. Total RNA was extracted using TRIzol reagent (Invitrogen; Thermo Fisher Scientific, Inc.), from three replicates of each *Srf* genotype. The quality and integrity of the RNA samples were examined as previously ^26^. RNA-seq was performed with Illumina Hiseq 2500 platform that included quality control, library preparation, fragmentation and PCR enrichment of target RNA according to standardized procedures. 150 bp paired-end raw reads were initially processed to obtain clean reads by removing adaptor sequences, low quality sequences and empty reads. After quality control, the clean reads were mapped to mouse genome (mm10) using STAT. Genes expression level were quantitated with TEtranscripts. Differentially expressed genes (DEGs) were identified with DESeq2. An absolute log2 fold change >1 and the adjusted p-value (FDR) significance score <0.05 were used as thresholds to identify DEGs. Gene set enrichment analysis (GSEA) was performed using GSEA software (**http://software.broadinstitute.org/gsea/**) with default parameters.

### Single-cell RNA-seq library preparation, sequencing and data analysis

10x Genomics Chromium platform was used to capture and barcode cells to generate single-cell Gel Beads-in-Emulsion (GEMs) by following the manufacturer’s protocol. The resulting libraries were sequenced on an Illumina NovaSeq 6000 System. Sequences from scRNA-seq were processed with Cellranger (v.2.0.0) software (10x Genomics). In short, demultiplexing, UMI (unique molecular identifier) collapsing and alignment to the mouse (Mus musculus) reference genome (version: mm10) was performed. Only confidently mapped, non-PCR duplicates with valid barcodes and unique molecular identifiers were used to generate the gene-barcode matrix that contained 954 cells for WT and 1073 cell for *Srf* KO iPSCs. Further analysis—including data cleaning, normalization, scaling quality and filtering was performed using the Seurat v3.1.1 ^47^. To exclude genes that might be random noise, we filtered genes whose expression was detected in fewer than 10 cells. To exclude poor quality cells that might result from multiplets or other technical noise, we filtered cells that were considered outliers (> third quartile + 1.5× interquartile range or < first quartile - 1.5× interquartile range) based on the number of expressed genes detected, the sum of UMI counts and the proportion of mitochondrial genes. In addition, we limited the proportion of mitochondrial genes to a maximum of 0.1 to further remove potential poor-quality data from broken cells. After removing unwanted cells from the dataset, we normalized the data by the total expression, multiplied by a scale factor of 10,000 and log-transformed the result. To reduce noise that may be introduced by considering all the genes, we used “FindVariableFeatures” function to select ~3,000 highly variable genes (HVGs) that contribute greatly to cell-to-cell variation. Then, the expression of the HVGs was used as the feature set for linear dimensional reduction of the data through principal component analysis (PCA) using Seurat’s RunPCA function. Uniform Manifold Approximation and Projection (UMAP) dimensional reduction was performed on the scaled matrix (with most variable genes only) using the first 30 components of principal component analysis (PCA) to obtain a two-dimensional representation of the cell states.

## Data availability

Sequencing data generated for this study have been deposited in GEO under accession code GSE210074. All data analyzed during this study are included in the manuscript and supporting files. Source data files are provided for all figures and figure supplements.

## Acknowledgement

Research reported in this publication was supported in part by DP2GM123507 (SG) and the Kutnick Family Foundation.

## Author Contributions

J.Z. performed all bioinformatics analysis, produced all figures and tables, and wrote the manuscript. S.G. supervised the project, designed experiments and wrote the manuscript. X.H. performed all cell culture experiments. X.H., Q.W. and S.G. provided additional technical and intellectual support. All the authors reviewed the manuscript.

## Figure legends

**Supplementary Figure 1.** Characteristics of scRNA-seq with density plot of number of expressed genes per cell. 960 and 857 transcriptomes (filtered and normalized with Seurat) from *Srf KO* and WT iPSCs, respectively, were analyzed.

**Supplementary Figure 2.** *Srf* KO iPSCs and CAF-1 KD cells share overlapping geneexpression patterns.

**(A)** Venn diagram showing the comparison between differentially expressed genes in *Srf* KO iPSCs and p150-depleted ES.

**(B)** Venn diagram showing the comparison between differentially expressed genes in *Srf* KO iPSCs and p60-depleted ES.

Genes with an FDR-adjusted *p* value ≤0.05 and a log_2_ fold change of at least 1 were considered to be differentially expressed.

**Supplementary Tables 1.** Differentially expressed genes in *Srf* KO iPSCs compared with WT.

**Supplementary Tables 2.** Mean gene expression, CV^2^ and Fano factor of all 5943 genes in *Srf* KO and WT iPSCs based on scRNA-seq.

## References

1 Angelidis, I, Simon, LM, Fernandez, IE, Strunz, M, Mayr, CH, Greiffo, FR et al. An atlas of the aging lung mapped by single cell transcriptomics and deep tissue proteomics. Nat Commun 2019 10, 963.

2 Bahar, R, Hartmann, CH, Rodriguez, KA, Denny, AD, Busuttil, RA, Dolle, ME et al. Increased cell-to-cell variation in gene expression in ageing mouse heart. Nature 2006 441, 1011.

3 Herndon, LA, Schmeissner, PJ, Dudaronek, JM, Brown, PA, Listner, KM, Sakano, Y et al. Stochastic and genetic factors influence tissue-specific decline in ageing C. elegans. Nature 2002 419, 808.

4 Martinez-Jimenez, CP, Eling, N, Chen, HC, Vallejos, CA, Kolodziejczyk, AA, Connor, F et al. Aging increases cell-to-cell transcriptional variability upon immune stimulation. Science 2017 355, 1433.

5 Desai, RV, Chen, X, Martin, B, Chaturvedi, S, Hwang, DW, Li, W et al. A DNA repair pathway can regulate transcriptional noise to promote cell fate transitions. Science 2021 373.

6 Torres-Padilla, ME & Chambers, I. Transcription factor heterogeneity in pluripotent stem cells: a stochastic advantage. Development 2014 141, 2173.

7 Choi, J, Huebner, AJ, Clement, K, Walsh, RM, Savol, A, Lin, K et al. Prolonged Mek1/2 suppression impairs the developmental potential of embryonic stem cells. Nature 2017 548, 219.

8 Chen, X, Burkhardt, DB, Hartman, AA, Hu, X, Eastman, AE, Sun, C et al. MLL-AF9 initiates transformation from fast-proliferating myeloid progenitors. Nat Commun 2019 10, 5767.

9 Cheung, P, Vallania, F, Warsinske, HC, Donato, M, Schaffert, S, Chang, SE et al. Single-Cell Chromatin Modification Profiling Reveals Increased Epigenetic Variations with Aging. Cell 2018 173, 1385.

10 Mahmoudi, S, Mancini, E, Xu, L, Moore, A, Jahanbani, F, Hebestreit, K et al. Heterogeneity in old fibroblasts is linked to variability in reprogramming and wound healing. Nature 2019 574, 553.

11 Izgi, H, Han, D, Isildak, U, Huang, S, Kocabiyik, E, Khaitovich, P et al. Inter-tissue convergence of gene expression during ageing suggests age-related loss of tissue and cellular identity. Elife 2022 11.

12 Lahoute, C, Sotiropoulos, A, Favier, M, Guillet-Deniau, I, Charvet, C, Ferry, A et al. Premature aging in skeletal muscle lacking serum response factor. PLoS One 2008 3, e3910.

13 Iram, T, Kern, F, Kaur, A, Myneni, S, Morningstar, AR, Shin, H et al. Young CSF restores oligodendrogenesis and memory in aged mice via Fgf17. Nature 2022 605, 509.

14 Connelly, JT, Gautrot, JE, Trappmann, B, Tan, DW, Donati, G, Huck, WT et al. Actin and serum response factor transduce physical cues from the microenvironment to regulate epidermal stem cell fate decisions. Nat Cell Biol 2010 12, 711.

15 Medjkane, S, Perez-Sanchez, C, Gaggioli, C, Sahai, E & Treisman, R. Myocardin-related transcription factors and SRF are required for cytoskeletal dynamics and experimental metastasis. Nat Cell Biol 2009 11, 257.

16 Posern, G & Treisman, R. Actin’ together: serum response factor, its cofactors and the link to signal transduction. Trends Cell Biol 2006 16, 588.

17 Schratt, G, Weinhold, B, Lundberg, AS, Schuck, S, Berger, J, Schwarz, H et al. Serum response factor is required for immediate-early gene activation yet is dispensable for proliferation of embryonic stem cells. Mol Cell Biol 2001 21, 2933.

18 Battich, N, Stoeger, T & Pelkmans, L. Control of Transcript Variability in Single Mammalian Cells. Cell 2015 163, 1596.

19 Esnault, C, Stewart, A, Gualdrini, F, East, P, Horswell, S, Matthews, N et al. Rho-actin signaling to the MRTF coactivators dominates the immediate transcriptional response to serum in fibroblasts. Genes Dev 2014 28, 943.

20 Gerber, A, Esnault, C, Aubert, G, Treisman, R, Pralong, F & Schibler, U. Blood-borne circadian signal stimulates daily oscillations in actin dynamics and SRF activity. Cell 2013 152, 492.

21 Li, S, Czubryt, MP, McAnally, J, Bassel-Duby, R, Richardson, JA, Wiebel, FF et al. Requirement for serum response factor for skeletal muscle growth and maturation revealed by tissue-specific gene deletion in mice. Proc Natl Acad Sci U S A 2005 102, 1082.

22 Ragu, C, Elain, G, Mylonas, E, Ottolenghi, C, Cagnard, N, Daegelen, D et al. The transcription factor Srf regulates hematopoietic stem cell adhesion. Blood 2010 116, 4464.

23 Costello, P, Sargent, M, Maurice, D, Esnault, C, Foster, K, Anjos-Afonso, F et al. MRTF-SRF signaling is required for seeding of HSC/Ps in bone marrow during development. Blood 2015 125, 1244.

24 Miano, JM, Ramanan, N, Georger, MA, de Mesy Bentley, KL, Emerson, RL, Balza, RO, Jr. et al. Restricted inactivation of serum response factor to the cardiovascular system. Proc Natl Acad Sci US A 2004 101, 17132.

25 Lamm, N, Ben-David, U, Golan-Lev, T, Storchova, Z, Benvenisty, N & Kerem, B. Genomic Instability in Human Pluripotent Stem Cells Arises from Replicative Stress and Chromosome Condensation Defects. Cell Stem Cell 2016 18, 253.

26 Hu, X, Liu, ZZ, Chen, X, Schulz, VP, Kumar, A, Hartman, AA et al. MKL1-actin pathway restricts chromatin accessibility and prevents mature pluripotency activation. Nat Commun 2019 10, 1695.

27 Kolodziejczyk, AA, Kim, JK, Tsang, JC, Ilicic, T, Henriksson, J, Natarajan, KN et al. Single Cell RNA-Sequencing of Pluripotent States Unlocks Modular Transcriptional Variation. Cell Stem Cell 2015 17, 471.

28 Nichols, J & Smith, A. Naive and primed pluripotent states. Cell Stem Cell 2009 4, 487.

29 Ying, QL, Wray, J, Nichols, J, Batlle-Morera, L, Doble, B, Woodgett, J et al. The ground state of embryonic stem cell self-renewal. Nature 2008 453, 519.

30 Macfarlan, TS, Gifford, WD, Driscoll, S, Lettieri, K, Rowe, HM, Bonanomi, D et al. Embryonic stem cell potency fluctuates with endogenous retrovirus activity. Nature 2012 487, 57.

31 Falco, G, Lee, SL, Stanghellini, I, Bassey, UC, Hamatani, T & Ko, MS. Zscan4: a novel gene expressed exclusively in late 2-cell embryos and embryonic stem cells. Dev Biol 2007 307, 539.

32 Zalzman, M, Falco, G, Sharova, LV, Nishiyama, A, Thomas, M, Lee, SL et al. Zscan4 regulates telomere elongation and genomic stability in ES cells. Nature 2010 464, 858.

33 Ishiuchi, T, Enriquez-Gasca, R, Mizutani, E, Boskovic, A, Ziegler-Birling, C, Rodriguez-Terrones, D et al. Early embryonic-like cells are induced by downregulating replication-dependent chromatin assembly. Nat Struct Mol Biol 2015 22, 662.

34 Hendrickson, PG, Dorais, JA, Grow, EJ, Whiddon, JL, Lim, JW, Wike, CL et al. Conserved roles of mouse DUX and human DUX4 in activating cleavage-stage genes and MERVL/HERVL retrotransposons. Nat Genet 2017 49, 925.

35 Whiddon, JL, Langford, AT, Wong, CJ, Zhong, JW & Tapscott, SJ. Conservation and innovation in the DUX4-family gene network. Nat Genet 2017 49, 935.

36 Arsenian, S, Weinhold, B, Oelgeschlager, M, Ruther, U & Nordheim, A. Serum response factor is essential for mesoderm formation during mouse embryogenesis. EMBO J 1998 17, 6289.

37 Akiyama, T, Xin, L, Oda, M, Sharov, AA, Amano, M, Piao, Y et al. Transient bursts of Zscan4 expression are accompanied by the rapid derepression of heterochromatin in mouse embryonic stem cells. DNA Res 2015 22, 307.

38 Eckersley-Maslin, MA, Svensson, V, Krueger, C, Stubbs, TM, Giehr, P, Krueger, F et al. MERVL/Zscan4 Network Activation Results in Transient Genome-wide DNA Demethylation of mESCs. Cell Rep 2016 17, 179.

39 Zhang, Y, Huang, Y, Dong, Y, Liu, X, Kou, X, Zhao, Y et al. Unique Patterns of H3K4me3 and H3K27me3 in 2-Cell-like Embryonic Stem Cells. Stem Cell Reports 2021 16, 458.

40 Maldonado, M, Luu, RJ, Ramos, ME & Nam, J. ROCK inhibitor primes human induced pluripotent stem cells to selectively differentiate towards mesendodermal lineage via epithelial-mesenchymal transition-like modulation. Stem Cell Res 2016 17, 222.

41 Rajan, A, Tien, AC, Haueter, CM, Schulze, KL & Bellen, HJ. The Arp2/3 complex and WASp are required for apical trafficking of Delta into microvilli during cell fate specification of sensory organ precursors. Nat Cell Biol 2009 11, 815.

42 Ikeda, T, Hikichi, T, Miura, H, Shibata, H, Mitsunaga, K, Yamada, Y et al. Srf destabilizes cellular identity by suppressing cell-type-specific gene expression programs. Nat Commun 2018 9, 1387.

43 Chang, J, Wei, L, Otani, T, Youker, KA, Entman, ML & Schwartz, RJ. Inhibitory cardiac transcription factor, SRF-N, is generated by caspase 3 cleavage in human heart failure and attenuated by ventricular unloading. Circulation 2003 108, 407.

44 Davis, FJ, Gupta, M, Pogwizd, SM, Bacha, E, Jeevanandam, V & Gupta, MP. Increased expression of alternatively spliced dominant-negative isoform of SRF in human failing hearts. Am J Physiol Heart Circ Physiol 2002 282, H1521.

45 Patten, LC, Belaguli, NS, Baek, MJ, Fagan, SP, Awad, SS & Berger, DH. Serum response factor is alternatively spliced in human colon cancer. J Surg Res 2004 121, 92.

46 Zhang, X, Azhar, G, Huang, C, Cui, C, Zhong, Y, Huck, S et al. Alternative splicing and nonsense-mediated mRNA decay regulate gene expression of serum response factor. Gene 2007 400, 131.

47 Macosko, EZ, Basu, A, Satija, R, Nemesh, J, Shekhar, K, Goldman, M et al. Highly Parallel Genome-wide Expression Profiling of Individual Cells Using Nanoliter Droplets. Cell 2015 161, 1202.

